# A novel method for estimating LDC levels: total cholesterol and non-HDLC can be used to predict the normal LDLC level in apparently healthy population

**DOI:** 10.1101/228130

**Authors:** Guo-Ming Zhang, Shumei Bai, Gao-Ming Zhang, Xiao-Bo Ma

## Abstract

**Background:** Triglycerides (TG), total cholesterol (TC), high-density lipoprotein cholesterol (HDLC), and low-density lipoprotein cholesterol (LDLC) are usually ordered together to test lipid metabolism in physical examination and clinical application. LDLC is a component of TC and its change is closely correlated to TC.

**Objective:** We aim to predict the normal LDLC level by using TG, TC, HDLC and non-HDLC (nonHDLC) in this study.

**Methods:** TG, TC, HDLC, and LDLC data were obtained from Laboratory Information System (LIS) based on 4 years (Octo, 1, 2013-Sept, 30, 2017) period health check-up TG, TC, HDLC, and LDLC(direct clearance method) were measured using TBA2000FR biochemical analyzer. The nonHDLC was calculated with TC minus HDLC. Correlation between TG, TC, nonHDLC, and LDLC were analyzed using Spearman’s rank approach. Receiver operating characteristics curve analysis was used to evaluate the predictive of TG, TC and nonHDLC for the normal LDLC level(less than 130mg/dL).

**Results:** Both TC(r = 0.870) and nonHDLC (r = 0.893) were significantly positively correlated with LDLC. Area under curve of TC and nonHDLC can be used to predict normal LDLC level. Optimal thresholds were 182.5 mg/dL(4.72 mmol/L) for TC and 135.3 mg/dL(3.50 mmol/L) for nonHDLC. Based on these optimal thresholds, less than 0.5% and 0.4% of tests with elevated LDLC might be missed, but the missing elevated LDLC is not too high(less than 147.3 mg/dL).

**Conclusion:** If the nonHDLC is less than 135.3mg/dL(3.50 mmol/L) and/or TC is less than 182.5mg/dL(4.72 mmol/L) for the apparently healthy populations the LDLC level will be less than 130mg/dL(3.36 mmol/L). Total cholesterol and non-HDLC can be used to predict the normal LDLC level in apparently healthy population is a novel method for estimating LDC level.

## INTRODUCTION

Triglycerides (TG), total cholesterol (TC), high-density lipoprotein cholesterol (HDLC), and Low-density lipoprotein cholesterol (LDLC) is commonly used in clinical practice [1]. The TC test is a measure of the level of total amount of cholesterol for a person, including LDLC and high-density lipoprotein cholesterol (HDLC) [2], as well as a certain type of triglycerides. The test of cholesterol is very important because elevated level of cholesterol can clog the arteries even lead to heart diseases and strokes [3]. Usually, serum TG, TC, HDLC, and LDLC are ordered together in the clinical laboratory. Since LDLC is a component of the TC and the change of LDLC is closely correlated to TC in most cases, we are wondering that LDLC may not be necessarily tested until the TC and nonHDLC are lower than the normal level. Thus, we proposed that serum normal LDLC level could be predicted by using TC and non-HDLC (nonHDLC) as reflex tests. Reflex testing was defined as automatically adding or removing a test by the biochemical analyzer for saving costs of clinical laboratories. For example, serum total bilirubin (TBIL) and conjugated bilirubin (CBIL) are highly correlated and approximately 87% of CBIL test reduced from TBIL test [4]. Most labs use automated technology for doing lipid profiles with automatic calculation of LDL-C by Friedewald in Europe and America, and all LDLC levels were measured using a direct clearance method on auto biochemical analyzer in China. However, the problem with Friedewald equation is that LDLC levels are commonly discordant with both direct clearance method, and the direct clearance method due to slightly higher costs. LDLC may not accurately reflect the true risk of LDL particles[3], the elevated LDLC should be measured by direct method. Hence, we search for a novel method for estimating the LDLC level. In this study, we analyzed the correlation between TG, TC, nonHDLC, and LDLC and concluded that the nonHDLC is under 135.3mg/dL(3.50 mmol/L) and/or TC is under 182.5mg/dL(4.72 mmol/L), LDLC is not more than 130mg/dL(3.36 mmol/L) for the apparently healthy population.

## MATERIALS AND METHODS

### Study cohort and data extraction

The data of TG, TC, HDLC and LDLC were obtained from laboratory information system (LIS). There are 34270 subjects (21651 males and 12619 females) (Table 1.) for health check-ups from October 2013 to September 2017 in Shuyang People’s Hospital were examined. Due to no complaint for visiting Shuyang People’s Hospital, we regard them as apparently healthy individuals. Age, gender, fasting serum TG (GPO-PAP no correction), TC (Cholesterol Oxidase method), HDLC (Direct Clearance Method) and LDLC (Direct Clearance Method) were extracted from LIS. TG, TC, HDLC, and LDLC were measured using TBA2000FR biochemical analyzer (Toshiba Co., Ltd., Japan). Regular quality control procedures is conducted every day in the laboratory medicine of Shuyang People’s Hospital. The external quality assessment scheme of Jiangsu Center for Clinical Laboratories were performed twice a year to validate the quality of TC, HDLC, and LDLC results. The ethics committee of ShuyangPeople’s Hospital approved this study.

**Table 1.** The characteristics of the participants

| Parameters | female | male | 2 |
| --- | --- | --- | --- |
|  | 12619 | 21651 | 3 |
| Age | 35 | 43 |  |
|  | (30-48) | (31-55) |  |
| Total Cholesterol(mmol/L) | 172.9 | 179.0 |  |
|  | (152.0-197.2) | (156.6-203.0) |  |
| Triglycerides(mmol/L) | 89.4 | 123.1 |  |
|  | (64.6-131.9) | (86.8-179.8) |  |
| LDLC(mmol/L) | 98.6 | 106.7 |  |
|  | (80.8-119.1) | (88.6-126.1) |  |
| nonHDLc(mmol/L) | 120.7 | 133.0 |  |
|  | ( 100.2-144.6 ) | ( 111.4-156.2 |  |
| HDLc(mmol/L) | 50.3 | 44.5 |  |
|  | (43.3-58.0) | (38.7-51.8) |  |

All data is represented bymedian and interquartile range.

### Statistical Analysis and Calculation

The nonHDLC was calculated from TC minus HDLC [1] and their relationship was analyzed using Spearman’s approach. For normal LDLC level, receiver operating characteristics curve analysis was used to evaluate the predictive accuracy of TG, TC, HDLC, and nonHDLC. Normal LDLC level were less than 130 mg/dL (3.36 mmol/L) [5, 6]. All statistical analyses were performed using EXCEL2007 (Microsoft Corporation, Beijing, China) and MedCalc 15.2.2 (MedCalc Software, Ostend, Belgium). A p value less than 0.05 was considered statistically significant.

## RESULTS

### The characteristics of the subjects

The data of 34270 pairs of cholesterols and triglycerides tests between October 1, 2013, and September 30, 2017 were extracted from the health examination center of Shuyang People’s Hospital, including 21651 males and 12619 females (Table 1).

### Correlation between TG, TC, HDLC, and nonHDLC and LDLC

Both TC (r = 0.870, *P* < 0.0001) and nonHDLC (r = 0.893, *P* < 0.0001) were significantly positively correlated with LDLC (**Figure 1**).

**Figure 1.**
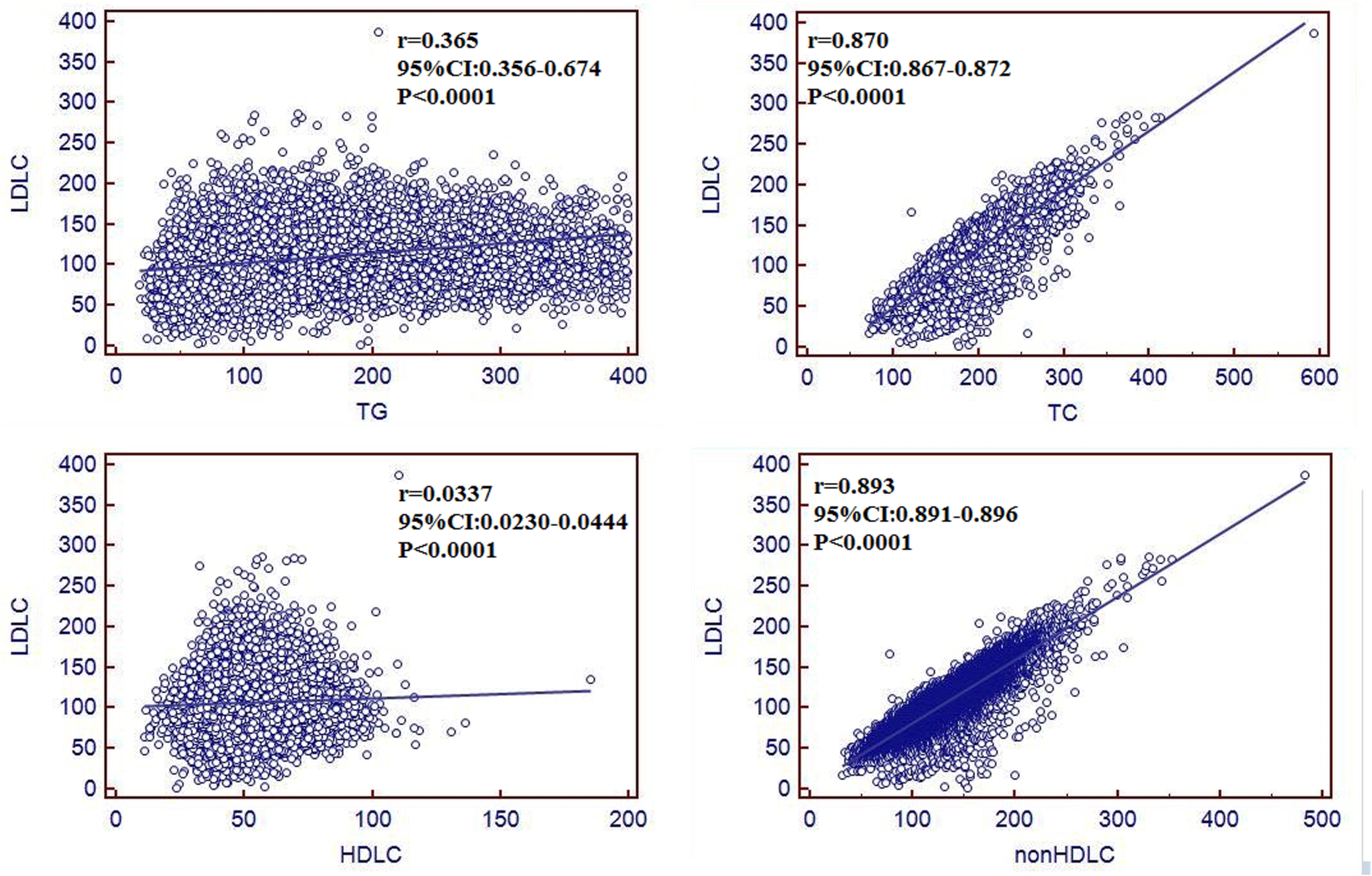
Scatter plots for low-density lipoprotein cholesterol (LDLC) and Triglycerides(TG), LDLC and total cholesterol(TC), LDLC and high-density lipoprotein cholesterol (HDLC) LDLC and non-density lipoprotein cholesterol (nonHDLC). Their relationship was analyzed using Spearman’s approach.

### The optimal threshold of TC and nonHDLC in predicting normal LDLC level

**Figure 2** shows the ROC curves of TG, TC, HDLC, and nonHDLC for predicting normal LDLC level (less than 130mg/dL). Areas under curve (AUC) were 0.675, 0.950, 0.541, and 0.957 for TG, TC, HDLC, and nonHDLC, respectively. As list in **Table 2**, nonHDLC was notably better than TC for predicting of normal LDLC level in terms of diagnostic performance and leakage in different TG levels. At these thresholds of TC and nonHDLC, less than 2.6% and 1.8% of tests with elevated LDLC might be missed but the missing elevated LDLC is lower(less than 147.3 mg/dL=3.81 mmol/L). As list in **Table 3**, when the nonHDLC is less than 135.3mg/dL(3.50 mmol/L) and/or TC is less than 182.5mg/dL(4.72 mmol/L), the LDLC will be less than 130mg/dL for the all the populations. If the nonHDLC is under 139.2mg/dL(3.60 mmol/L) and/or TC is under 182.5mg/dL(4.72 mmol/L), the LDLC will be low 130mg/dL for the populations (The TG is less than 400mg/dL).

**Figure 2.**
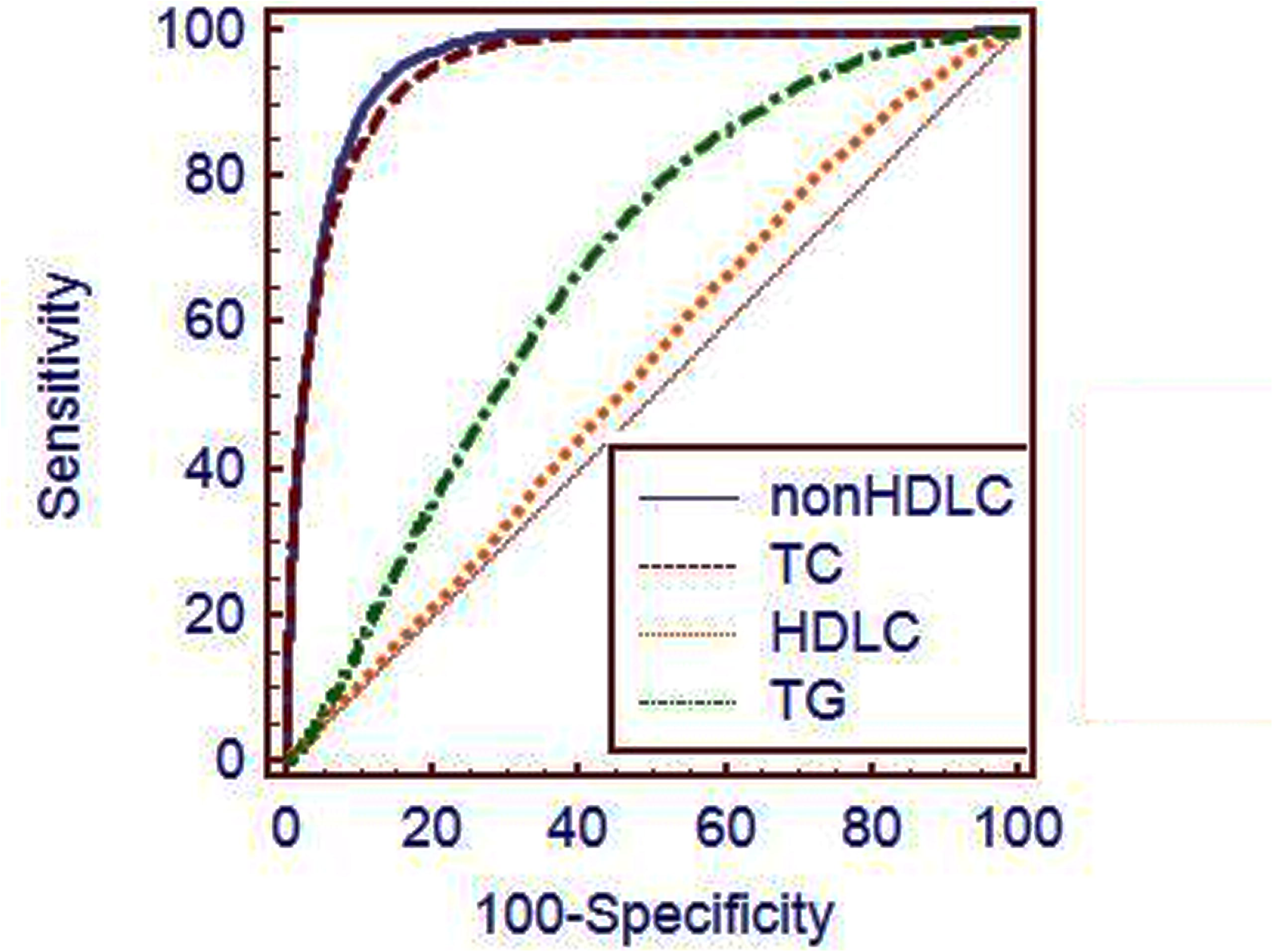
Receiver operating characteristics curves of Triglycerides(TG), total cholesterol (TC), high-density lipoprotein cholesterol (HDLC), and low-density lipoprotein cholesterol (LDLC) for predicting normal low-density lipoprotein cholesterol(LDLC).

**Table 2.**
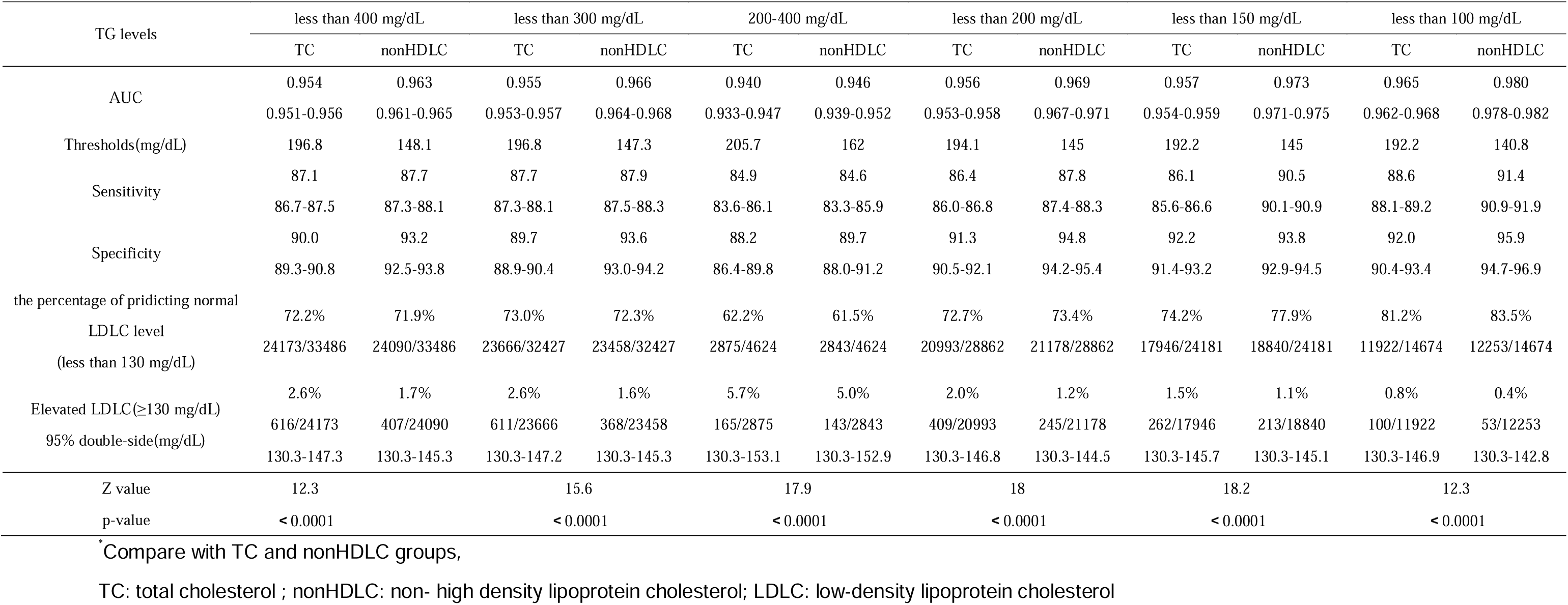
The optimal threshold and accuracy of total cholesterol and non-high density lipoprotein cholesterol, and its performance in predicting normal low-density lipoprotein cholesterol

**Table 3.** The optimal threshold total cholesterol and non-high density lipoprotein cholesterol, and their performance in predicting normal 1ow-density lipoprotein cholesterol

| TG levels | total |  | less than 400 mg/dL |  | 200-400 mg/dL |  | less than 200 mg/dL |  |
| --- | --- | --- | --- | --- | --- | --- | --- | --- |
|  | TC | nonHDLc | TC | nonHDLc | TC | nonHDLc | TC | nonHDLc |
| The optimal thresholds | 182.5 | 135.3 | 182.5 | 139.2 | 186.8 | 147.7 | 181.4 | 138.4 |
| The percentage of predicting<br>normal LDLc level<br>(less than 130 mg/dL) | 56.0% | 61.7% | 57.3% | 62.7% | 40.7% | 44.2% | 59.4% | 66.5% |
|  | 19175/34270 | 21138/34270 | 19175/33486 | 20995/33486 | 1884/4624 | 2043/4624 | 17132/28862 | 19206/28862 |
| Elevated LDLc($\geq 130$ mg/dL) | 0.5% | 0.4% | 0.5 | 0.4% | 0.6 | 0.5% | 0.4 | 0.3% |
|  | 87/19175 | 83/21138 | 87/19175 | 83/20995 | 11/1884 | 11/2043 | 71/17132 | 65/19206 |
| 95% double-side(mg/dL) | 130.3-147.3 | 130.3-145.3 | 130.3-147.3 | 130.3-145.3 | 130.3-139.6 | 130.3-138.8 | 130.3-147.3 | 130.3-145.3 |

TC: total cholesterol ; nonHDLc: non- high density lipoprotein cholesterol; LDLc: low-density lipoprotein cholesterol

## DISCUSSION

In this study, we analyzed serum TG, TC, nonHDLC, LDLC and their correlation in a large cohort of apparently healthy subjects, and we found that TC and nonHDLC were positively correlated with LDLC. Subjects have low TC and nonHDLC levels usually accompany with a low LDLC levels and vice versa. The results of ROC for predicting normal LDLC level has supported our hypothesis. The AUC was more than 0.95 for both TC and nonHDLC, indicating that TC and nonHDLC have accuracy for predicting normal LDLC level. Therefore, serum lower LDLC levels(less than 130mg/dL) could be predicted by using TC and non-HDLC. Given the diagnostic performance and the proportion of elevated LDLC, nonHDLC is notably better than TC for predicting normal LDLC level. Two optimal thresholds of TC and nonHDLC for predicting normal LDLC level were 182.5 mg/dL(4.72 mmol/L) and 139.2 mg/dL(3.60 mmol/L) (TG is less than 400mg/dL). If TC is less than 182.5 mg/dL and/or nonHDLC is less than 139.2 mg/dL, there will be not more than 130 mg/dL(3.36 mmol/L) for LDLC levels. According to these thresholds, less than 0.5% and 0.4% of elevated LDLC could be missed, and the missing elevated LDLC is lower (Table 2 and Table 3). In this study, approximately 56% of LDLC test or calculation could be reduced. The price of a LDLC testing is approximately 4RMB (0.65 USD) [4] in China, thus, the cost for LDLC test would be greatly saved. If nonHDLC is used as a reflex test for LDLC (when elevated LDLC is more than 130 mg/dL), approximately 11.3 million LDLC test would be reduced and 7.2 million USD would be saved annually. The direct LDLC test will automatically be performed when the predicting LDLC level is not normal level. This saves time for the doctor by not needing to order the direct LDLC test, saves time for the patient by not needing to draw another blood sample, and speeds up the time to obtain the accurate LDLC results. Compared with the Friedewald formula [7] and the formula is considered inaccurate(TG level is more than 220mg/dL)[8], in this study we take into account the cost savings but also screen out the normal LDLC level, and it was not affected by triglyceride levels.

There are some limitations in this study. First, this is a single-center study. Second, the subjects in this study come from health check-ups. Third, if the LDLC level is less than 130 mg/dL, there will be less than 0.5% for missing elevated LDLC.

The results of present study indicate that a large number of normal LDLC level can be predicted by using TC and/or nonHDLC. To our knowledge, this study is the first to predict LDLC level by using receiver operating characteristics curve analysis.The approach of this study may be suited for other subjects. However, because of difference of the detection system and subjects, they should get the optimal threshold based on local data for using TC and/or nonHDLC to predict normal LDLC level.

## Acknowledgements

None

